# A statistical-based method for the construction and analysis of gene network: application to bacteria

**DOI:** 10.1101/2024.09.03.611021

**Authors:** Zhiyuan Zhang, Guozhong Chen, Erguang Li

## Abstract

Bacteria play a crucial role in environmental conservation, human health, and medicine. Whether in the gut or the soil, bacterial genomes are rich repositories of resources, such as exploring potential drugs and biopesticides. However, our ability to develop new therapies and deepen our understanding of the bacterial world is hindered by the largely unknown functions of bacterial genes. In this study, we proposed a method of gene network construction and analysis based on a Gaussian Graphical Model (GGM) and random sampling strategy to infer direct interactions at the genomic level in bacteria. Using *Vibrio cholerae* and *Staphylococcus aureus* as examples, we integrated partial correlation-based gene co-expression data with gene regulatory and essentiality information extracted from public databases to construct more comprehensive gene networks. Networks built upon bacterial different phenotypes, such as biofilm formation, flagellar assembly, and stress response, demonstrate the effectiveness of this method in deciphering unknown gene functions, uncovering new phenotype-associated factors, and identifying their corresponding interactions, thus providing new targets for experimental validation by researchers. Additionally, we extended this method to 14 bacteria, including 13 pathogens, supporting the investigation of gene functions and pathways at the genomic level in these bacteria. More importantly, for other species, this method of gene network construction can be easily implemented, provided that sufficient transcriptome sequencing samples are available.

## Introduction

The study of bacterial gene function and regulatory mechanism lags significantly behind equivalent research in human, progressing slowly (1,2). Currently, approximately 20% to 50% of genes in some bacteria remain unannotated, hindering breakthrough in fields such as antibiotic resistance and bacterial pathogenicity (3,4). Traditional methods for annotating bacterial gene function primarily rely on experimental approaches like gene knockout and overexpression, which are costly, time-consuming, and challenging to apply on genomic scale (5,6,7). This inevitably results in delays in bacterial gene function annotation and slow progress in studying gene regulatory mechanism.

Gene network has now been widely utilized to investigate gene regulatory mechanism and predict unknown gene function. They also serve as an effective tool for identifying functional relationship between genes. Numerous studies have found that the functional annotations (biological processes and pathways) represented by modules identified from gene network aid in understanding the functions of genes encoding hypothetical (uncharacterized) proteins (8,9,10). Gene network can help uncover gene regulatory mechanism, such as by identifying key regulatory factors and their associated genes during bacterial pathogenesis, thereby aiding in the elucidation of bacterial pathogenic mechanism (11). Some studies have used gene co-expression network to decipher the function of unannotated genes and the pathways they represent (12,13).

The construction of gene network depends on mathematical model designed to capture and describe the complex relationships and dynamic properties within gene expression data. Among these, pairwise correlation based on pearson correlation coefficient is a classic method for constructing gene co-expression network (14,15). However, this approach measures only marginal relationships between variables, making it unable to distinguish between direct and indirect effect. That is, if two genes are correlated, it does not necessarily mean they directly depend on each other, as the observed correlation may be mediated by a third gene. Another popular computational method is Weighted Gene Co-expression Network Analysis (WGCNA) (16,17,18,19), which essentially transforms Pearson correlation between gene pairs into weights, often by raising the correlation coefficient to a power. Network constructed by WGCNA can reflect the relationship between genes and identify gene modules associated with phenotype. However, this method still does not solve the problem of distinguishing direct from indirect correlation between gene pairs. Another approach, the Graphical Gaussian Model (GGM), uses partial correlation, measuring the residual correlation after accounting for the effect of other genes, to determine the correlation between gene pairs. This method has been used in several studies to construct gene network (20, 21). However, the classical GGM theory is inadequate for situations where the number of genes (p) is much greater than the number of samples (n), a common scenario in transcriptome experiments. Several improved methods, such as the least absolute shrinkage and selection operator (LASSO) regularization (22) and covariance shrinkage (23), have been proposed to address this “high-dimensional low-sample” problem by approximating sparse precision matrix estimation. Nevertheless, the introduction of optimization techniques cannot eliminate the excessive noise caused by the small sample size, which leads to the generation of gene networks with many false positive connections, making it difficult to capture true gene relationships.

Here, we developed a method based on the Gaussian Graphical Model (GGM) combined with random sampling for constructing and analyzing bacterial gene network. By introducing a mathematical model for random sampling, which calculates the number of sampling iterations *N*, we ensure that the number of genes in a single estimate is roughly equal to the number of samples, thus reducing the gap between p and n, causing reduction in noise. Additionally, we introduced the concept of multi-channels, where each channel independently estimates gene pair partial correlations. We have drawn inspiration from the concept of multi-channels in convolutional neural network, which is introduced to differentiate it from the number of iterations required for random sampling in this study, and essentially involves multiple independent estimates of grenome-wide gene pairs for partial correlation (pcor). Subsequently, we performed overlapping analysis on the gene pairs across different channels to identify shared gene pairs for network construction, thereby enhancing network stability. We also integrated prior knowledge, such as transcriptional regulatory information, gene essentiality, and protein interaction data, into the network, making it more comprehensive and systematic. Finally, we constructed and analyzed gene networks for 14 common bacteria and developed a gene network visualization platform, Bnetwork, to present the analyzed results. We believe that Bnetwork will greatly assist researchers in the field of bacteriology.

### Developed method for constructing bacterial gene network

In this study, we combined Gaussian Graphical Model (GGM) with a random sampling strategy to construct bacterial gene network. The detailed implementation process is illustrated in Figure 1. For collection and preprocessing, we obtained raw transcriptome sequencing data of bacteria from the EBI ENA (24) and NCBI SRA (25) databases. After data collection, we processed the transcriptome data, including the removal of non-compliant sequences using FastQC (26) and Trim_galore (27) and the mapping and transcript quantification using Salmon (28), ultimately obtaining TPM values. During this process, we assessed the sample quality based on the mapping rate, discarding samples with a mapping rate below 50%. The gene expression values of samples that met quality standard were concatenated according to experimental conditions, forming an overall gene expression matrix. To mitigate the influence of non-expressed genes on subsequent analysis, we used an in-house program to exclude genes that were not expressed in over 50% of the experimental conditions. Since the GGM requires gene expression data to follow a normal or skewed distribution, we performed a log transformation on the gene expression matrix to achieve requirement. After obtaining the log-transformed gene expression matrix, we used the ComBat function of Sva package in R to remove batch effects between different experimental conditions (29).

**Figure 1.**
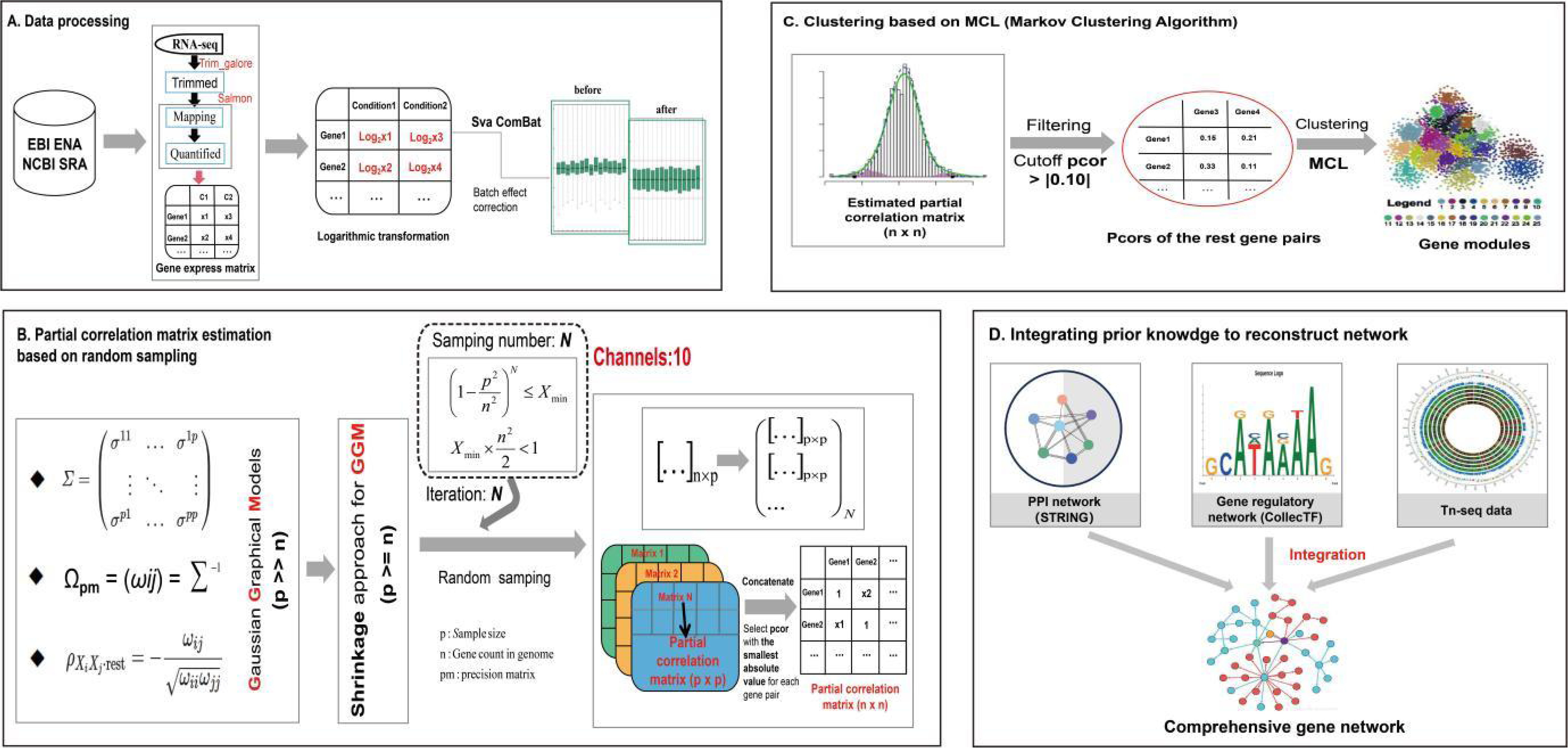
The workflow of statistical-based method

The pcors of gene pair derived from the GGM were used to measure the direct correlation between gene pairs. Classic GGM theory requires the setting for p >> n, but in practical applications, the number of next-generation sequencing samples (**p**) in bacteria is often much smaller than the number of corresponding bacterial genes (**n**). To address this issue, we employed the Shankage method (), a technique that incorporates regularization and moderation to infer GGM, allowing the **p** to be slightly greater than or even equal to the **n**. Additionally, we introduced a random sampling strategy, where p genes are selected each time, corresponding to the number of sequencing samples, without considering the order of gene pairs. In theory, we could obtain gene pairs 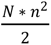 by repeating this sampling *N* times, ensuring that the number of selected gene pairs fully covers genome-wide gene pairs. We constructed the corresponding mathematical model and derived the times of sampling iteration *N*.

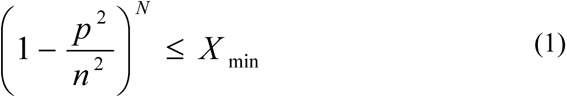

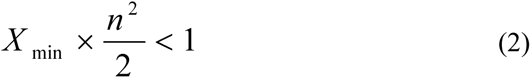

In this context, p represents the number of samples, n represents the number of species genes, *N* represents the times of sampling iteration, 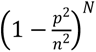 represents the probability of a gene pair not being sampled in *N* iterations, and 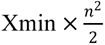 represents the minimum acceptable number of gene pairs not covered at the genomic level, which we set to be less than 1 in this study. This indicates that we aim to theoretically cover all gene pairs at the genomic level. By deriving, we calculated the number of sampling iterations *N*.

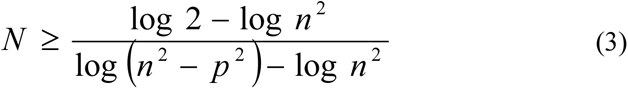

During the random sampling process, we estimated the partial correlations (pcors) among p genes after each sampling, ultimately obtaining *N* p×p partial correlation matrices. Each gene pair was theoretically sampled multiple times *C*, so multiple pcors were generated for each gene pair.

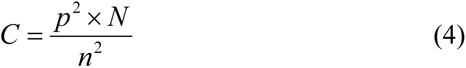

In this study, we selected the smallest absolute pcor as the final estimate for the extended network, representing the one with the largest amount of effects from other genes removed. After replacing each gene pair with its smallest absolute pcor, we obtained the pcors of genome-wide gene pairs resulting in an n×n partial correlation matrix.

Upon obtaining the pcors of genome-wide gene pair, we considered that biologically meaningful gene networks are sparse, meaning that most nodes in the network are not connected (pcor = 0). Unlike the LASSO method, which may increase the number of false negatives, the Shankage method does not provide a sparse estimate of the partial correlation matrix but instead uses a pre-thresholding approach to estimate partial correlations (pcors). Therefore, we set the threshold to an absolute pcor of 0.1 (20), considering gene pairs with an absolute pcor below 0.1 as having no correlation.

Multi-channels is introduced to this study, with 10 channels used to independently estimate the partial correlations (pcors) of gene pairs, resulting in 10 n × n partial correlation matrices for following analysis. We then defined the pcor threshold, calculated p-values for pcors, and corrected for multiple testing (30). Pcors were considered significant only when they exceeded the threshold, and were depicted as edges in network. After obtaining the qualified gene pairs, we performed overlapping analysis of the qualified gene pairs from the 10 channels, identifying shared gene pairs across multiple channels. These gene pairs and their corresponding pcors were extracted and subjected to clustering analysis using the Markov clustering algorithm (MCL) (31, 32), obtaining corresponding gene modules.

Finally, we integrated prior knowledge to guide the construction of the gene network. The primary sources of prior data included protein interaction data from the STRING database (33, 34), transcription factor regulatory information from databases like CollecTF (35), and well-processed Tn-seq data related to bacterial phenotype studies from OGEE (36). By integrating multiple sources of biological information, we aimed to construct a network that comprehensively reflects the complex relationships between genes and proteins, thereby helping researchers better understand the complexity and multi-level regulatory mechanisms of biological system.

### An illustration depicting the construction and analysis of gene network, exemplified by *Vibrio cholerae*

*Vibrio cholerae* is the primary pathogen responsible for cholera, an acute intestinal infectious disease (37, 38). Currently, with the rapid advancement of next-generation sequencing technologies, transcriptomic studies of *Vibrio cholerae* have generated a vast amount of data and information. These data will be utilized for gene network construction.

Initially, we downloaded RNA-seq samples of *Vibrio cholerae* from the ENA and SRA databases, totaling 705 samples (Figure 2A). We employed a standardized RNA-seq analysis pipeline, where the mapping step utilized the reference transcriptome of *Vibrio cholerae N16961*, comprising 3,912 genes. The TPM values of the genes were obtained through analysis. Subsequently, we filtered out genes not expressed under more than 50% of experimental conditions and samples with less than 50% of reads mapping to the reference transcriptome, discarding low-quality genes and samples. We obtained 360 high-quality samples and 3,852 normally expressed genes, resulting in a 360 × 3,852 gene expression matrix. Next, we performed log transformation on the gene expression matrix. After obtaining the log-transformed gene expression matrix, we removed batch effects between different experiments using the Sva ComBat function in R (Figure 2C).

**Figure 2.**
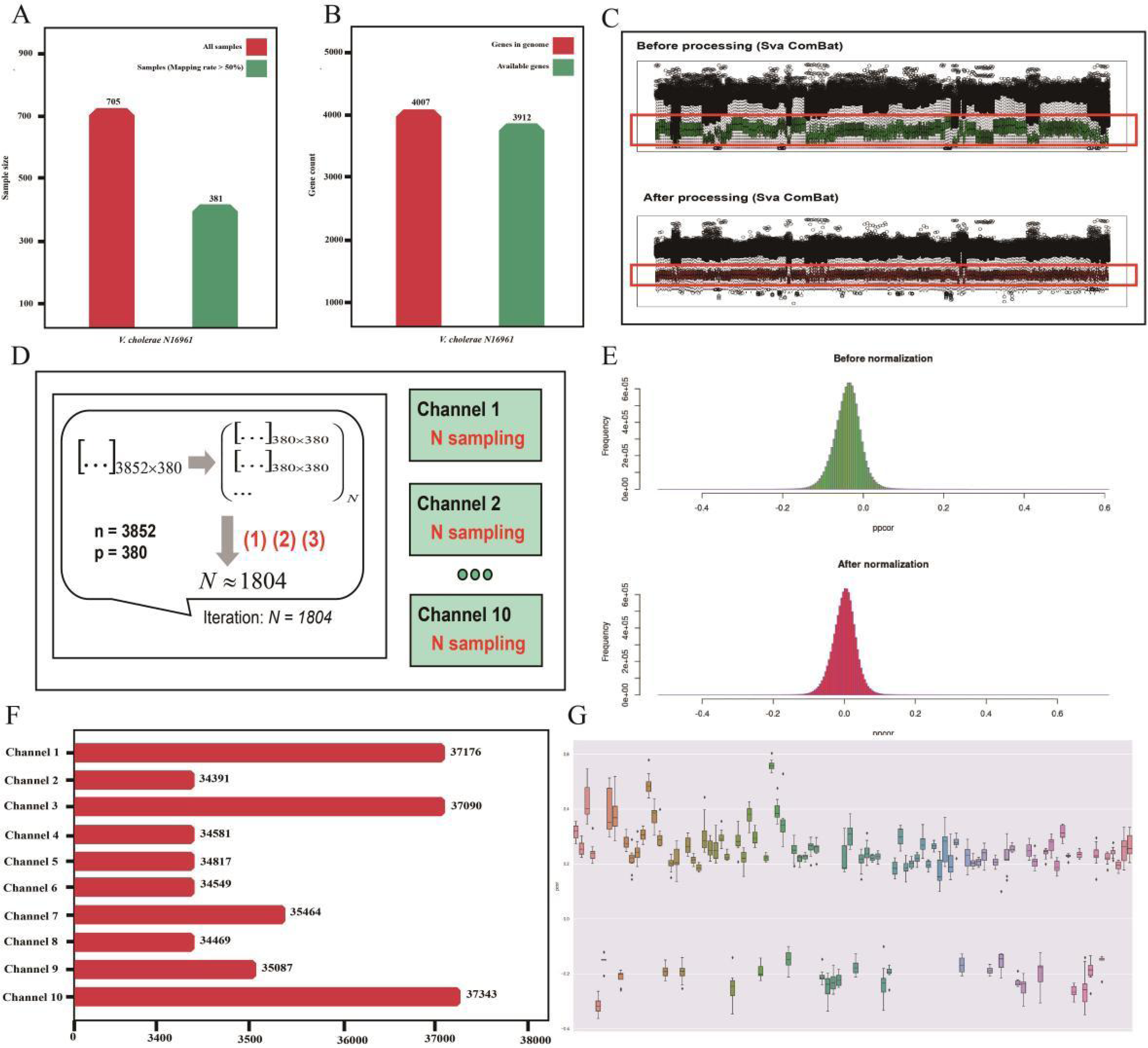
Details of workflow using *Vibrio cholerae* as an example

Subsequently, based on the sample size of 360, we needed to randomly select 360 genes from the 3,852 genes, repeating this process ***N*** times to fully cover the gene pairs at the genomic level. We calculated *N* = 1,804 using formulas (1), (2), and (3). Due to the 10 channels, the pcor estimate of genome-wide gene paris were required to independently repeat 10 times. During the computation of the partial correlation matrix, the GeneNet package (39) was employed to estimate the pcor for each gene pair, using the Shankage method to infer the GGM. After computation, we performed Fisher’s z-transformation on the estimated pcor values to enhance accuracy (40). As shown in Figure 2E, most gene pairs had pcors close to 0, indicating a lack of interaction, consistent with previous studies suggesting that connections in biomolecular networks are generally sparse. Setting a threshold of 0.1 for the absolute value of pcor, we discarded all gene pairs below this threshold for all channels.

Following the above procedure, we estimated the pcors for gene pairs within each channel. According to the statistics, each channel obtained between 34,000 and 38,000 significant gene pairs, which would serve as nodes in the gene network, with the corresponding pcors representing the edges. In this study, the multi-channel approach was employed to improve the accuracy of pcor estimates for gene pairs. By performing 10 independent pcor estimates, the minimum pcor was taken as the final pcor for the gene pairs, reducing the impact of other genes and noise on the results. Through a boxplot analysis, we found slight differences in the pcors of 100 shared gene pairs randomly extracted from the 10 channels, although no contradictory pcors were observed across channels. Integrating pcors across multiple channels proved to be an effective method for reducing variability. It should be noted that increasing times of sampling directly could achieve similar results. However, the multi-channel approach significantly reduces the computational load by filtering gene pairs within each channel before integrating the pcors across channels, thus lowering computational costs.

The specific integration process involved overlapping analysis of gene pairs filtered in channel 1 with those in channel 2, retaining shared gene pairs with the smallest absolute pcors. This process continued sequentially through channel 10, ultimately obtaining shared gene pairs for subsequent clustering analysis and network construction. To assess the impact of the number of channels on network construction variability, we calculated the number and proportion of gene pairs discarded during each overlap analysis. As shown in Figure 3B, the number of discarded gene pairs was initially large, but as the overlapping analysis progressed, the number and proportion of discarded gene pairs began to decrease significantly, eventually stabilizing. In the final three analysis, the number and proportion of discarded gene pairs were 611 (5.0%), 448 (3.8%), and 352 (3.1%), respectively, with a small gap. Through this analysis, we obtained 11,240 usable gene pairs, involving 3,520 *Vibrio cholerae* genes.

**Figure 3.**
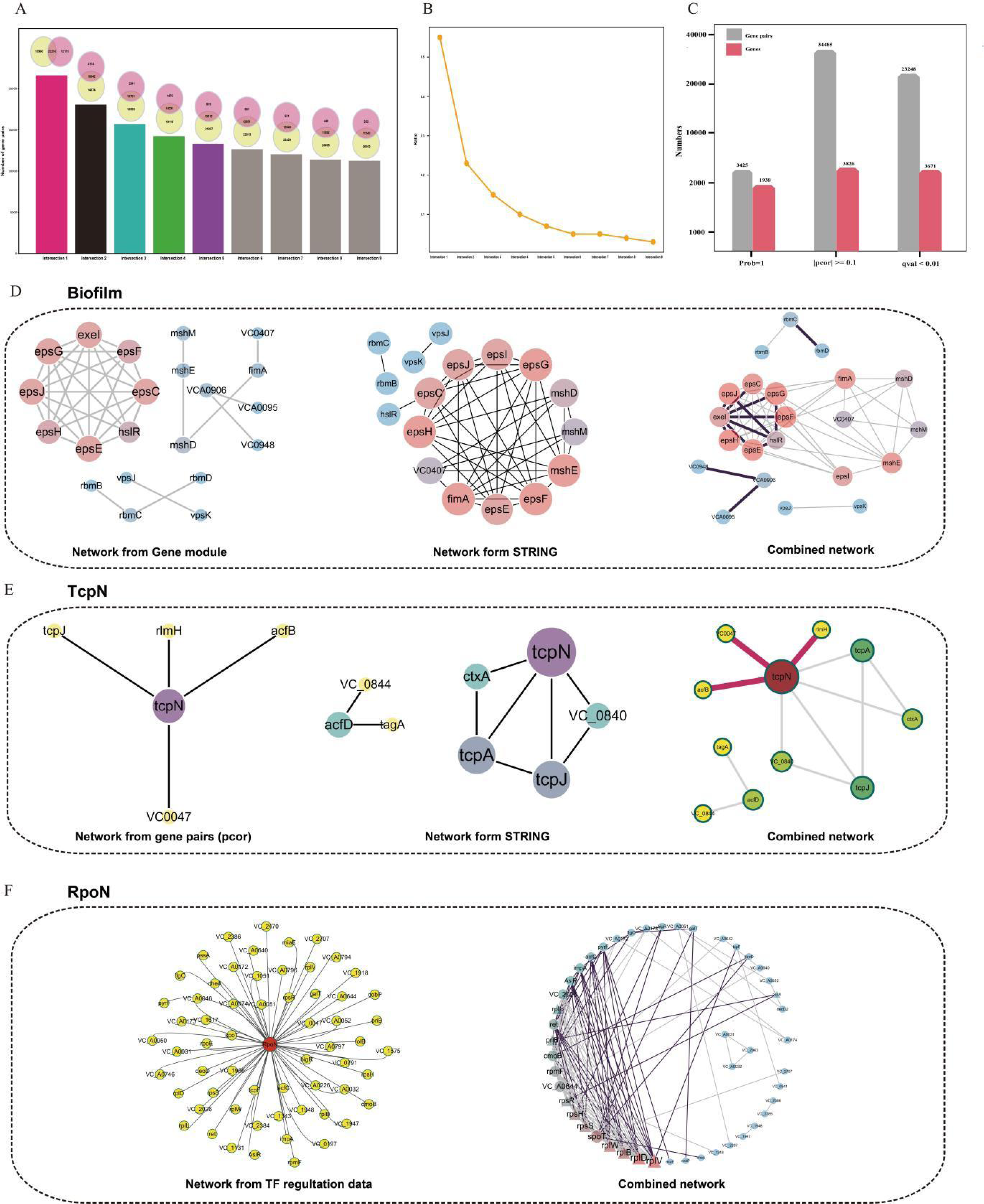
An introduction of various gene network types

Additionally, we also explored utilizing probability derived from model and q-value adjusted for multiple testing corrections to filter for significant gene pairs, which resulted in a smaller gene pair scale compared to the pcor threshold-based method, indicating a more stringent gene network. We then performed clustering analysis on the retained 11,240 gene pairs using the MCL algorithm with parameters “–I 1.2 – scheme 7”, obtaining 270 gene modules, 177 of which contained more than 5 gene members. We analyzed these modules, finding that up to 170 of them exhibited interactions among genes, and 100 modules were enriched in GO (41) and KEGG (42) pathways (Table S2). These results indicate that our proposed method corresponds well with existing experimental research findings while also providing novel, unexplored genes and related interactions for further investigation.

We integrated the pcor-based gene co-expression network with prior knowledge, including protein (gene) interaction information from the STRING database, transcriptional regulatory information from the collecTF database, and gene essentiality information from the OGEE database (36) obtained through Tn-seq data analysis, to obtain a more comprehensive *Vibrio cholerae* gene network.

Using the biofilm phenotype, as well as the TcpN and RpoN transcription factors as examples, we introduced different types of gene networks formed through the integration of various data. Firstly, an integrated analysis of gene module information with STRING data for biofilm phenotype of *Vibrio cholerae*. Through GO and KEGG functional enrichment analysis, we identified gene modules enriched in biofilm-related pathways. One such module, containing 21 gene members, includes eight genes *espJ*, *espE*, *espC*, *espG*, *espH*, *espF*, *VC0411*, and *VC0405* that are involved in biofilm formation (43, 44). We then extracted pcors for these gene pairs to construct an initial network, resulting in 36 gene pairs. Simultaneously, we retrieved interaction data from the STRING database for these genes in the module, which are often derived from experimental data. Finally, we integrated these datasets to construct a hybrid gene network comprising 56 gene pairs, closely associated with biofilm formation. The hybrid network includes several novel, unannotated genes (Figure 3D), captured by the network based on pcor, such as *VCA0095*, which is connected to membrane-associated genes *VCA0906* and *VC0948*. These genes may be involved in the biofilm formation of *Vibrio cholerae*. Additionally, there are connections between biofilm-associated genes and genes of unknown function, marked as edges in the figure, suggesting potential interactions between them.

The pcor data of the *tcpN* gene, which encodes the TcpN transcription factor, was combined with transcriptional regulation and protein interaction information to construct gene network. Firstly, genes correlated with *tcpN* were identified in the pcors, including *tcpJ*, *VC0007*, *rlmH*, and *acfB*. Subsequently, we extracted information on genes regulated by the TcpN transcription factor from public databases such as CollecTF and Mr.Vc (14) (Figure S1), and inputted TcpN and its regulated genes into the STRING database to obtain their interaction data. These pieces of information were then integrated to visualize the gene network. Correlations were observed among the gene pairs *tcpN*-*VC0007* (pcor, 0.1153), *tcpN*-*rlmH* (pcor, 0.1164), and *tcpN*-*acfB* (pcor, 0.3532), which were not captured by public databases. These results suggest that the *VC0007*, *rlmH*, and *acfB* genes may interact with TcpN at the protein level and might even be regulated by the TcpN transcription factor. However, further experimental validation is required to determine the specific regulatory and interaction mechanism.

Based on the known RpoN transcriptional regulatory network, which includes 61 regulated genes (14, 45), the gene network was constructed by integrating pcors and STRING database protein (gene) interaction data (Figure S2). Since RpoN is involved in ribosome synthesis, we analyzed the essentiality of RpoN and its regulated genes using Tn-seq information from OGEE database. We discovered that 7 genes in this regulatory network are essential, highlighting the indispensable and non-redundant nature of ribosome synthesis in *Vibrio cholerae*. Finally, all data were integrated and visualized, clearly demonstrating that multiple interactions between genes were captured in the hybrid network formed by our developed method. This finding underscores the superiority of our approach in uncovering gene functions and potential interactions.

### Comparison with existing studies

We compared our findings with other studies that constructed *Vibrio cholerae* gene networks using different methods. Two studies employed WGCNA for network construction (13, 18), a method requiring a low number of sequencing samples and generally, more than ten samples are sufficient for analysis. However, a limited sample size inevitably introduces noise, and WGCNA cannot exclude indirect correlations between genes (16), leading to many false-positive gene pairs. As shown in Figure 4A, the three studies identified 6, 49, and 177 gene modules (more than 5 gene members), respectively. This result is expected, as WGCNA cannot distinguish between direct and indirect correlations, resulting in many false-positive gene pairs and clustering unrelated genes into the same module. From these studies, we extracted the final number of gene pairs obtained. DuPai’s study ultimately identified more then 1,000,000 gene pairs, while our study identified 11,400 gene pairs (Figure 4B). It is worth noting that Qin’s study did not provide weighted gene co-expression data, so corresponding gene pair data could not be obtained. Additionally, we extracted gene *viuD*, an ABC transporter with the most connections from the 11,400 gene pairs, in total connections with 49 genes. When analyzed using DuPai’s constructed weighted gene co-expression network, *viuD* was found to be correlated with 448 genes, far exceeding the number of connections obtained through GGM method. In the STRING interaction data, only 18 genes were found to interact with *viuD,* among which 3 genes were shared with the pcor-based network, and 4 were shared with the WGCNA-based network, suggesting that the WGCNA method generated some false-positives lacking biological significance.

**Figure 4.**
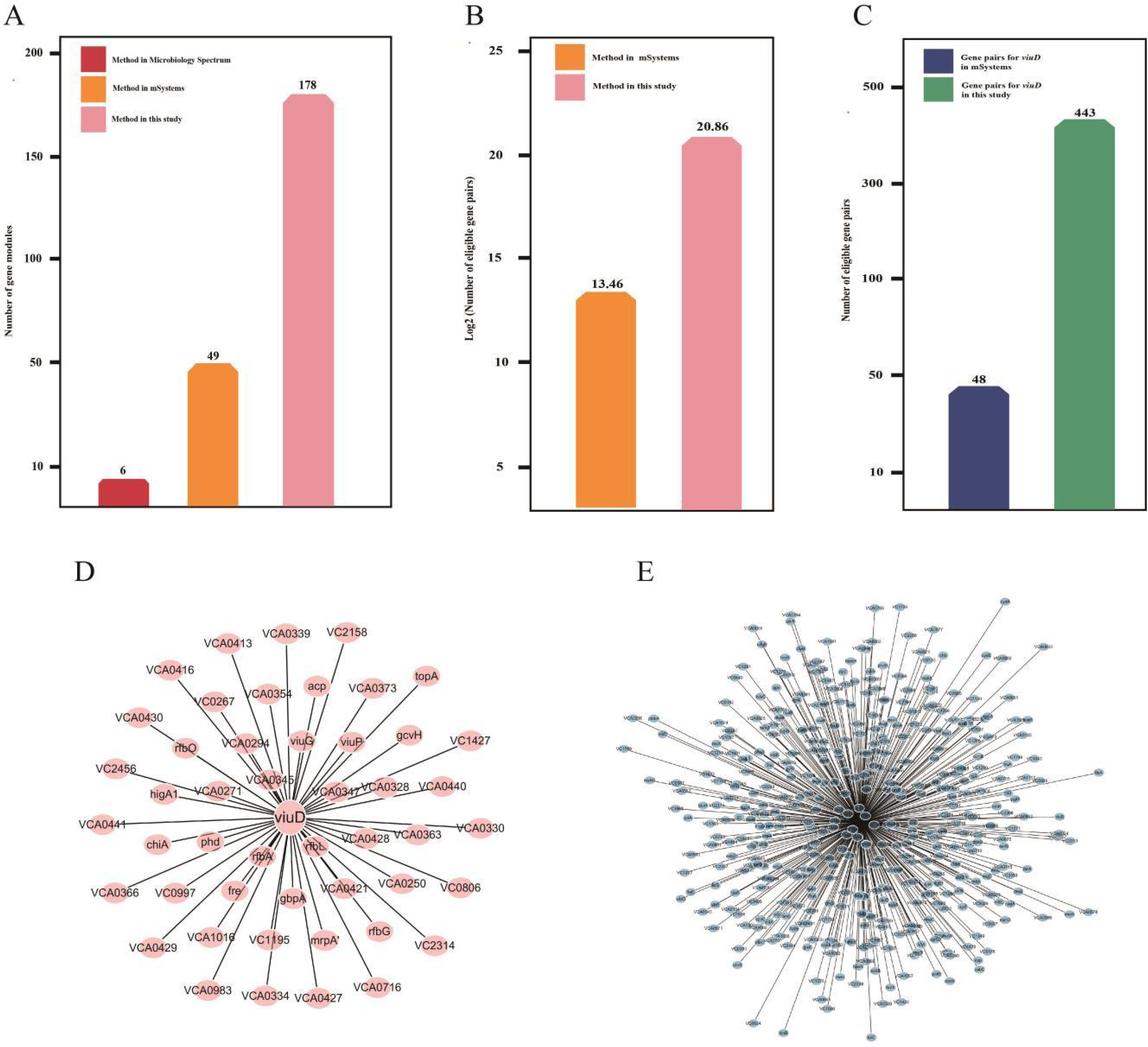
A comparison with other studies

**Figure 5.**
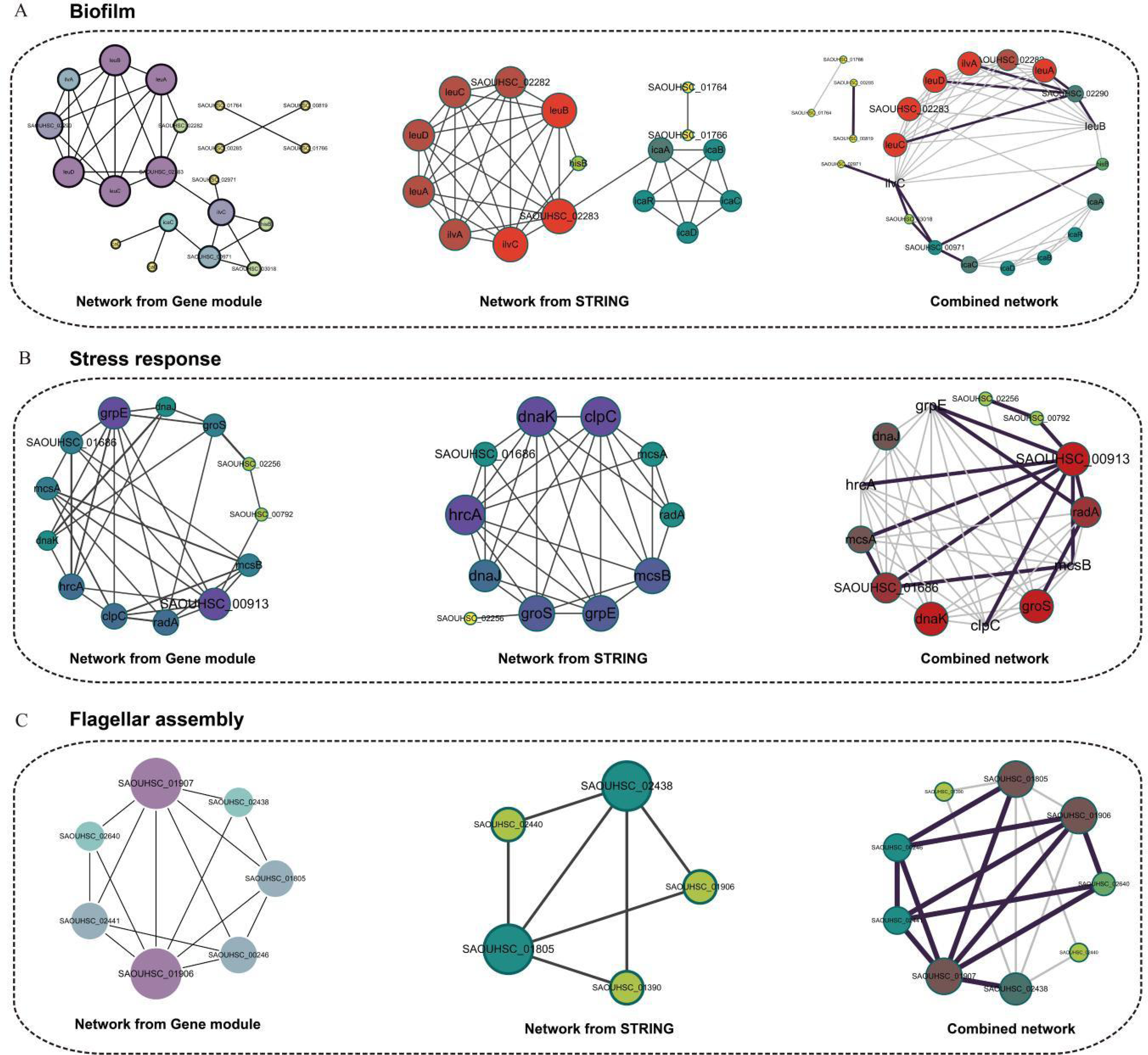
Analysis of three phenotypic networks in *Staphylococcus aureus*

### Analysis of phenotypic network in *Staphylococcus aureus*

We explored gene network in *Staphylococcus aureus* under three phenotypic conditions, including biofilm formation, stress response, and flagellar assembly. For the biofilm phenotype of *Staphylococcus aureus*, current public databases, including GO (Gene Ontology) and KEGG (Kyoto Encyclopedia of Genes and Genomes), lack annotations for the biofilm formation pathway. Therefore, we sourced relevant information through literature review. We identified that genes associated with biofilm formation in *Staphylococcus aureus*, such as *icaA*, *icaB*, *icaC*, *icaD*, and *icaR*, which produce the intercellular polysaccharide adhesin controlling biofilm formation (46, 47), and *hisB* (48), known for mutations affecting biofilm formation, cluster within the same gene module, suggesting a close association with biofilm formation. GO and KEGG analysis of this module revealed enrichment in the amino acid synthesis pathway (FDR: 7.46e-06).

We extracted gene pairs within this module based on estimated pcors and used these gene pairs to construct network. Additionally, we compared this network with an interaction network based on the STRING database. There were 21 shared gene pairs between the two networks, including interactions among *icaB*, *icaC*, and *icaD* genes, which have been experimentally validated, indicating that the pcor-based network aligns with experimental findings (49). Furthermore, our network revealed additional edges representing previously unrecognized interactions, such as the *SAOUHSC_00971*-*hisB* and *SAOUHSC_00971*-*icaC* gene pairs. *SAOUHSC_00971*, an uncharacterized protein, shows correlation with known biofilm-associated genes, suggesting it may play a significant role in the biofilm formation, which is consistent with the analysis of biofilm-associated protein predictor (50). Notably, the gene network derived from the STRING database integrates not only gene co-expression information but also protein-protein interactions, such as those between proteins encoded by SAOUHSC_02282 and SAOUHSC_02283. Thus, combining these two networks into a hybrid network provides more comprehensive information, guiding further experimental research.

Through GO and KEGG analysis of these gene modules, we discovered that in one module containing 13 genes, 8 genes including *grpE*, *dnaJ*, *dnaK*, *clpC*, *hrcA*, *radA*, *mcsB*, and *mcsA* were enriched in the stress response pathway (21 genes) (FDR: 9.53E-12). Therefore, we hypothesize that this gene module is associated with the bacterial stress response, with the remaining 5 genes possibly participating in the stress response. To verify this hypothesis, we analyzed *SAOUHSC_01686,* coding chaperone protein, and its interactions with 4 known stress response genes including *grpE*, *dnaJ*, *dnaK*, and *hrcA*. *SAOUHSC_02256*, predicted as an abortive infection protein, has GO analysis suggesting involvement in membrane processes, while STRING results indicate its correlation with GroES and GroEL. GroES, together with the chaperonin GroEL, plays a crucial role in assisting protein folding and responding to environmental stress (51). *SAOUHSC_00792* is a coding conserved hypothetical protein, and *SAOUHSC_00913*, a conserved hypothetical protein, belong to the LysR transcriptional regulatory family involved in biological regulation. We extracted gene pairs from this module based on pcors and constructed corresponding network. It was observed that SAOUHSC_00913, showing a high similarity with the transcriptional regulator LysR, connects with 8 genes in the module, indicating a relationship. However, no interactions between *SAOUHSC_00913* and other genes in the module were found in the STRING database. Considering SAOUHSC_00913 as an upstream regulatory factor, it should regulate multiple downstream genes theoretically. The hybrid network shows that *SAOUHSC_00913* interacts with several genes involved in stress response, implying that SAOUHSC_00913 may play a crucial role in stress response. The additional edges in the hybrid network provide new directions for studying the regulatory mechanism of the stress response in *Staphylococcus aureus*.

GO and KEGG analysis of the gene modules revealed that in one module containing 9 genes, and 5 genes, including *SAOUHSC_01906*, *SAOUHSC_01805*, *SAOUHSC_02438*, *SAOUHSC_01390*, and *SAOUHSC_02440* were enriched in the flagellar assembly pathway (5 of 18 genes, FDR: 2.85e-07). Interaction data extracted from the STRING database showed no interactions among the remaining 4 genes and other genes in the module. However, the pcor-based network indicated connections between genes such as *SAOUHSC_01907*, *SAOUHSC_00246*, *SAOUHSC_02441*, and *SAOUHSC_02640* with genes involved in flagellar assembly, highlighting the superiority of our developed method in uncovering gene interactions. By integrating these data, we redrew the network, revealing multiple additional edges surpassing the known edges from public databases, indicating that many gene interactions remain to be explored.

### Development of the Bnetwork Analysis Platform

We have also employed this method to construct networks for 11 other common human pathogens, including *Acinetobacter baumannii*, *Pseudomonas aeruginosa*, *Salmonella enterica*, *Streptococcus pneumoniae*, *Mycobacterium tuberculosis*, *Streptococcus pyogenes*, *Listeria monocytogenes*, *Porphyromonas gingivalis*, *Enterococcus faecalis*, *Klebsiella pneumoniae*, *Streptococcus mutans*, and the soil microorganism *Bacillus subtilis*. As shown in Figure 6A, the number of samples for each bacterium, the number of genes involved, and the times of random sampling are all presented. The process of network construction generated vast amounts of data, such as the pcors of gene pair, gene modules, their corresponding functional enrichment information, gene annotations, and transcriptional regulation data, all of which are of significance. To ensure researchers in related fields can access and understand these data, we have developed an analysis platform (https://bnetwork.biownmcli.info/#!/), named Bnetwork. This platform provides gene annotation and network information, along with their visualizations, for 14 bacteria. Additionally, transcriptome data, gene regulation information, and gene essentiality data analyzed in this study can be directly downloaded from Bnetwork. We hope Bnetwork can serve as a foundational infrastructure for the bacterial research community.

**Figure 6.**
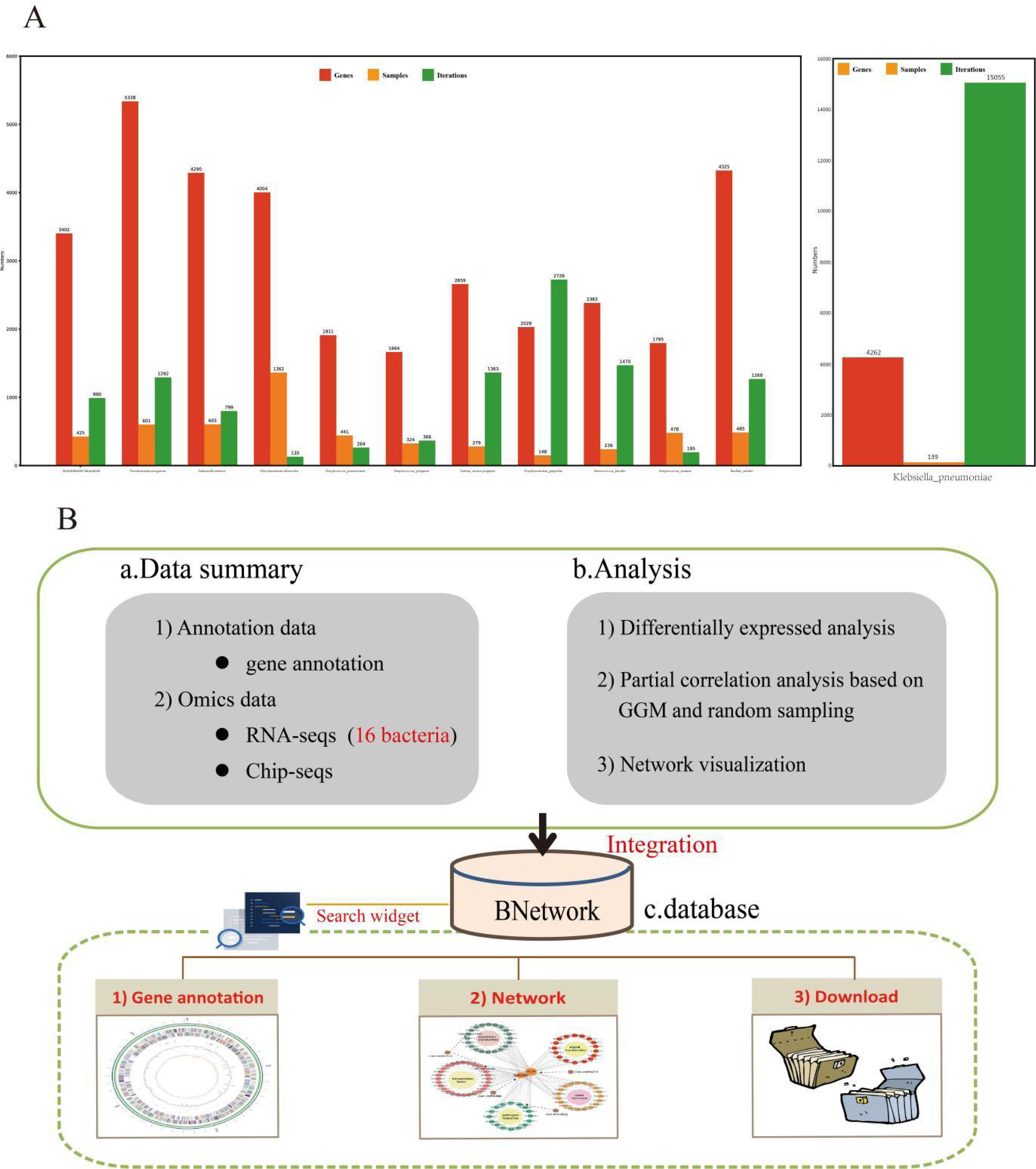
Introduction of Bnetwork platform

## Discussion

The study of gene network is of profound significance in understanding the functional mechanism of biological systems. Presently, gene network mapping relies primarily on experimental methods, such as inferring regulatory relationships through gene knockout and knockdown, confirming protein-protein interactions through co-immunoprecipitation (52), and computational approaches, such as constructing co-expression networks by calculating pearson correlation coefficients between gene pairs from gene expression data. Although experimental methods provide direct gene interaction data, they have limitations, such as high costs, low throughput, and the inability to generate vast amounts of gene interaction data in a short period (5, 6, 7, 53). In contrast, computational methods offer significant cost savings and can generate gene networks quickly, while also integrating data from public databases like STRING, OGEE to construct more comprehensive network. However, it must be noted that combining computational methods with experimental validation remains the best strategy for constructing accurate and reliable gene network.

In this study, we developed a method for constructing bacterial gene network based on GGM and pcors that is also applicable to other species, allowing the accurate identification of direct interactions between genes and filtering out indirect interactions, which have a straightforward interpretation as conditional independencies, distinguishing direct from indirect effects. Prior to this research, networks constructed based on pearson correlation or WGCNA inevitably introduced a large number of false positive connections, which obscured biologically meaningful networks and hindered researchers from uncovering valuable gene interactions. For examples, in the comparison with existing research mentioned above, we discovered that when using the ViuD interaction information from the STRING database as a reference, where ViuD interacts with 18 proteins, however, the number of connections for *viuD* (VC0777) in the network constructed based on WGCNA was 448, indicating associations with 448 genes. Among these, 4 genes were shared with the WGCNA network (4/18, 22.2%). In contrast, the GGM-based method of network construction connected *viuD* to 49 genes, with 3 genes in shared with the WGCNA network (3/18, 16.7%). The result appears that WGCNA is more efficient and accurate than the GGM in network construction. However, considering the accuracy of the detected gene pairs, the correctness rate of gene pairs in the network constructed based on the GGM (3/49) was 6.8 times higher than that based on WGCNA (4/448). Therefore, we believe that the WGCNA method may have produced some biologically meaningless false positives.

Additionally, when dealing with transcriptomic sequencing data, the high dimensionality, where the number of genes far exceeds the number of samples, inevitably leads to noise in the constructed gene network. To address this, we combined regularization, using a shrinkage approach in graphical Gaussian modeling, with random sampling, precisely determining the number of samples based on statistical model, an aspect often overlooked in other studies. This aimed to reduce noise while overcoming the p>>n limitation in classical GGM. Furthermore, multi-channels analysis was introduced, inspired by the VGG model. In this study, we independently estimated the pcors of genome-wide gene pairs across different channels, and performed overlapping analysis on gene pairs that met the criteria (|pcor| ≥ 0.1) across different channels. The shared gene pairs across channels were used to construct the gene network, thus removing unstable gene pairs and enhancing the network’s stability. However, it is important to note that when sample sizes are too small, noise is unavoidable regardless of optimization techniques (54). Increasing the sample size can address this issue. Single-cell transcriptomics is a technique used to study gene expression levels within individual cells (55, 56, 57). Using microdroplet technology, single-cell transcriptome sequencing experiments can sequence thousands to tens of thousands of cells simultaneously. We envision that if these cells are treated as samples, single-cell transcriptome sequencing could generate thousands or even tens of thousands of samples in a single experiment, leading to networks that are more stable and valuable compared to those constructed from traditional transcriptome sequencing samples. In future studies, we will explore the construction of gene networks using single-cell transcriptome data, despite the significant challenges posed by this approach, such as weak co-expression signals in single cells and the frequent presence of missing expression values in single-cell datasets (58, 59).

Lastly, recognizing the importance of prior knowledge in constructing systemic gene networks. We incorporated protein-protein interaction data from public databases such as STRING, transcriptional regulation data from CollecTF and Mr.Vc databases, and bacterial gene essentiality information from the OGEE database into the gene co-expression networks constructed based on GGM. Taking *Vibrio cholerae* and *Staphylococcus aureus* as examples, we constructed gene networks with gene modules as the core, integrating prior knowledge. In this process, we discovered that gene modules obtained through MCL were enriched in various biological pathways, such as biofilm formation, stress response, and ribosome synthesis. This indicates that certain gene members within the modules participate in specific pathways, while also suggesting that the remaining gene members may be involved in the same biological processes. This hybrid network can uncover new genes and interactions, significantly aiding researchers in deciphering the functions of unannotated genes and the multifunctionality of known genes.

Additionally, to facilitate the use of the data generated in this study by researchers, we developed a gene network visualization platform, Bnetwork, to display gene network information for 14 bacteria. We are confident that Bnetwork will provide substantial assistance to bacterial researchers.

## Materials and methods

### Transcriptome data collection and processing

For data collection, we collected and curated transcriptome datasets from 14 bacteria research of *Acinetobacter baumannii*, *Pseudomonas aeruginosa*, *Salmonella enterica*, *Streptococcus pneumoniae*, *Mycobacterium tuberculosis*, *Streptococcus pyogenes*, *Listeria monocytogenes*, *Porphyromonas gingivalis*, *Enterococcus faecalis*, *Klebsiella pneumoniae*, *Streptococcus mutans*, *Vibrio cholerae*, *Staphylococcus aureus* and *Bacillus subtilis*, including 6,750 RNA-seq samples, from the European Nucleotide Archive (EBI ENA, https://www.ebi.ac.uk/ena/browser/home) (24) and Sequence Read Archive (SRA, https://www.ncbi.nlm.nih.gov/sra/) (25). For the processing of raw sequencing reads, FastQC (http://www.bioinformatics.babraham.ac.uk/projects/fastqc/) was used to evaluate the overall quality of the raw sequencing reads, followed by the Trim_galore to remove sequencing vectors and low-quality bases (26) and performed transcript quantification using Salmon (28), which adopted TPM (Transcript Per Million) for normalization (Figure 1B), a better unit for RNA abundance than RPKM and FPKM since it respects the invariance property and is proportional to the average relative RNA molar concentration (60). It is essential to emphasize that the “mapping” step is based on salmon, which maps the reads obtained by next-generation sequencing onto the transcript of bacteria to achieve the estimation of gene expression. The batch effect correction between different experimental conditions is mainly performed by the ComBat function of sva package in R. In addition, all the transcriptomics are from Illumina sequencing, including single-end reads and pair-end reads. For pair-end reads, the use of FastQC and Trim_Galore requires the inspection and pruning of reads at both ends respectively, and the use of mapping tool (Salmon) need to compare read1 and read2 at the same time to ensure that the reads are positioned correctly. In order to make gene expression data follow normal or skewed distribution, A log transformation were performed on the gene expression matrix.

### Other data collection

We extracted protein-protein interaction information, which is primarily derived from experiments, data mining, gene co-expression, and other public repositories, from the STRING database (33). During the data extraction process, we set a high confidence of interaction score 0.7 as threshold, in order to filter out substandard protein-protein interactions. We downloaded gene essentiality information from the OGEE database (36), which analyzes Tn-seq data from bacteria to identify essential genes and genes associated with specific phenotypes. The CollecTF database compiles gene regulation data for bacteria, including regulatory factors, regulatory sites, regulatory mechanisms, and the corresponding regulated genes, aiding us in constructing a more systematic gene network (35). We gathered gene regulation information and gene annotation data for *Vibrio cholerae* from the Mr.Vc database (14).

### Network construction and visualization

After performing the pcor estimates of genome-wide gene pairs and filtering gene pairs using a pcor threshold of 0.1, the overall pcor-based gene network is depicted as shown (Figure S2). This network was arranged and visualized using Cytoscape software (61). For the hybrid network formed by the gene modules obtained through the MCL algorithm along with other biological information, we customized the network visualization by adjusting Cytoscape parameters to distinguish the core nodes within the network and to present potential edges.

### Database design

Bnetwork was designed as a relational database, with all data meticulously stored within a MySQL database. The website’s frontend was crafted using JavaScript and HTML, while the backend was developed using PHP, enhanced by the Slim framework to facilitate queries to the MySQL database and to offer RESTful APIs for programmable access to our data. The AngularJS framework was used to bride the front- and back-ends. For frontendan Apache server.

## Data availability

visualizations, Echarts.js and Plotly.js were utilized. The website is hosted on

Publicly available datasets were analyzed in this study. These data and corresponding code for the construction and analysis of gene network can be found here: https://bnetwork.biownmcli.info/#!/.

## Supporting information

vibrio cholerae

## Acknowledgements

The work was supported by grant from the National Key R&D Program of China (2023YFC2308200), Wannan Medical College key project research fund (WK2023ZZD08).

## References

1. Galperin MY, Koonin EV. From complete genome sequence to ’complete’ understanding? Trends Biotechnol. 2010;28(8):398–406.

2. Sorek R, Cossart P. Prokaryotic transcriptomics: a new view on regulation, physiology and pathogenicity. Nat Rev Genet. 2010;11(1):9–16.

3. Land M, Hauser L, Jun SR, Nookaew I, Leuze MR, Ahn TH, Karpinets T, Lund O, Kora G, Wassenaar T, Poudel S, Ussery DW. Insights from 20 years of bacterial genome sequencing. Funct Integr Genomics. 2015;15(2):141–61.

4. Price MN, Wetmore KM, Waters RJ, Callaghan M, Ray J, Liu H, Kuehl JV, Melnyk RA, Lamson JS, Suh Y, Carlson HK, Esquivel Z, Sadeeshkumar H, Chakraborty R, Zane GM, Rubin BE, Wall JD, Visel A, Bristow J, Blow MJ, Arkin AP, Deutschbauer AM. Mutant phenotypes for thousands of bacterial genes of unknown function. Nature. 2018;557(7706):503–509.

5. Hu P, Janga SC, Babu M, Díaz-Mejía JJ, Butland G, Yang W, Pogoutse O, Guo X, Phanse S, Wong P, Chandran S, Christopoulos C, Nazarians-Armavil A, Nasseri NK, Musso G, Ali M, Nazemof N, Eroukova V, Golshani A, Paccanaro A, Greenblatt JF, Moreno-Hagelsieb G, Emili A. Global functional atlas of Escherichia coli encompassing previously uncharacterized proteins. PLoS Biol. 2009;7(4):e96.

6. Deutschbauer A, Price MN, Wetmore KM, Tarjan DR, Xu Z, Shao W, Leon D, Arkin AP, Skerker JM. Towards an informative mutant phenotype for every bacterial gene. J Bacteriol. 2014;196(20):3643–55.

7. Nichols RJ, Sen S, Choo YJ, Beltrao P, Zietek M, Chaba R, Lee S, Kazmierczak KM, Lee KJ, Wong A, Shales M, Lovett S, Winkler ME, Krogan NJ, Typas A, Gross CA. Phenotypic landscape of a bacterial cell. Cell. 2011;144(1):143–56.

8. De R, Whiteley M, Azad RK. A gene network-driven approach to infer novel pathogenicity-associated genes: application to *Pseudomonas aeruginosa* PAO1. mSystems. 2023;8(6):e0047323.

9. Mao L, Van Hemert JL, Dash S, Dickerson JA. Arabidopsis gene co-expression network and its functional modules. BMC Bioinformatics. 2009;10:346.

10. Beiki H, Nejati-Javaremi A, Pakdel A, Masoudi-Nejad A, Hu ZL, Reecy JM. Large-scale gene co-expression network as a source of functional annotation for cattle genes. BMC Genomics. 2016;17(1):846.

11. Luo M, Chen G, Yi C, Xue B, Yang X, Ma Y, Qin Z, Yan J, Liu X, Liu Z. Dps-dependent in vivo mutation enhances long-term host adaptation in *Vibrio cholerae*. PLoS Pathog. 2023;19(3):e1011250.

12. Rosa BA, Jasmer DP, Mitreva M. Genome-wide tissue-specific gene expression, co-expression and regulation of co-expressed genes in adult nematode Ascaris suum. PLoS Negl Trop Dis. 2014;8(2):e2678.

13. DuPai CD, Wilke CO, Davies BW. A Comprehensive Coexpression Network Analysis in *Vibrio cholerae*. mSystems. 2020;5(4):e00550–20.

14. Zhang Z, Chen G, Hussain W, Qin Z, Liu J, Su Y, Zhang H, Ye M. Mr.Vc v2: An updated version of database with increased data of transcriptome and experimental validated interactions. Front Microbiol. 2022;13:1047259.

15. Cao D, Xu N, Chen Y, Zhang H, Li Y, Yuan Z. Construction of a Pearson- and MIC-Based Co-expression Network to Identify Potential Cancer Genes. Interdiscip Sci. 2022;14(1):245–257.

16. Langfelder P, Horvath S. WGCNA: an R package for weighted correlation network analysis. BMC Bioinformatics. 2008;9:559.

17. Xu M, Zhou H, Hu P, Pan Y, Wang S, Liu L, Liu X. Identification and validation of immune and oxidative stress-related diagnostic markers for diabetic nephropathy by WGCNA and machine learning. Front Immunol. 2023;14:1084531.

18. Qin ZX, Chen GZ, Yang QQ, Wu YJ, Sun CQ, Yang XM, Luo M, Yi CR, Zhu J, Chen WH, Liu Z. Cross-Platform Transcriptomic Data Integration, Profiling, and Mining in *Vibrio cholerae*. Microbiol Spectr. 2023;11(3):e0536922.

19. Zhang ZY, Chen GZ, Pan YY, Yang Z, Liu Y, Li EG. An in-depth analysis and exploration with focus on the biofilm in *Staphylococcus aureus*. bioRxiv 2024.05.05.592613; doi: 10.1101/2024.05.05.592613.

20. Ma S, Gong Q, Bohnert HJ. An Arabidopsis gene network based on the graphical Gaussian model. Genome Res. 2007;17(11):1614–25.

21. Xu Y, Wang Y, Ma S. SingleCellGGM enables gene expression program identification from single-cell transcriptomes and facilitates universal cell label transfer. Cell Rep Methods. 2024;4(7):100813.

22. Tibshirani R. Regression shrinkage and selection via the lasso. Journal of the Royal Statistical Society: Series B (Methodological). 1996; 58(1):267–288.

23. Schäfer J, Strimmer K. A shrinkage approach to large-scale covariance matrix estimation and implications for functional genomics. Stat Appl Genet Mol Biol. 2005;4:Article32.

24. Cummins C, Ahamed A, Aslam R, et al. The European Nucleotide Archive in 2021. Nucleic Acids Res. 2022;50(D1):D106–D110.

25. Sayers EW, O’Sullivan C, Karsch-Mizrachi I. Using GenBank and SRA. Methods Mol Biol. 2022;2443:1–25.

26. Utturkar S, Dassanayake A, Nagaraju S, Brown SD. Bacterial Differential Expression Analysis Methods. Methods Mol Biol. 2020;2096:89–112.

27. Fóthi Á, Liu H, Susztak K, Aranyi T. Improve-RRBS: a novel tool to correct the 3’ trimming of reduced representation sequencing reads. Bioinform Adv. 2024;4(1):vbae076.

28. Patro R, Duggal G, Love MI, Irizarry RA, Kingsford C. Salmon provides fast and bias-aware quantification of transcript expression. Nat Methods. 2017;14(4):417–419.

29. Castaldo R, Brancato V, Cavaliere C, et al. A Framework of Analysis to Facilitate the Harmonization of Multicenter Radiomic Features in Prostate Cancer. J Clin Med. 2022;12(1):140.

30. Schäfer J, Strimmer K. An empirical Bayes approach to inferring large-scale gene association networks. Bioinformatics. 2005;21(6):754–764.

31. Enright AJ, Van Dongen S, Ouzounis CA. An efficient algorithm for large-scale detection of protein families. Nucleic Acids Res. 2002;30(7):1575–1584.

32. Azad A, Pavlopoulos GA, Ouzounis CA, Kyrpides NC, Buluç A. HipMCL: a high-performance parallel implementation of the Markov clustering algorithm for large-scale networks. Nucleic Acids Res. 2018;46(6):e33.

33. Szklarczyk D, Kirsch R, Koutrouli M, et al. The STRING database in 2023: protein-protein association networks and functional enrichment analyses for any sequenced genome of interest. Nucleic Acids Res. 2023;51(D1):D638–D646.

34. Szklarczyk D, Gable AL, Lyon D, et al. STRING v11: protein-protein association networks with increased coverage, supporting functional discovery in genome-wide experimental datasets. Nucleic Acids Res. 2019;47(D1):D607–D613.

35. Kiliç S, White ER, Sagitova DM, Cornish JP, Erill I. CollecTF: a database of experimentally validated transcription factor-binding sites in Bacteria. Nucleic Acids Res. 2014;42(Database issue):D156–D160.

36. Chen WH, Minguez P, Lercher MJ, Bork P. OGEE: an online gene essentiality database. Nucleic Acids Res. 2012;40(Database issue):D901–D906.

37. Hsiao A, Zhu J. Pathogenicity and virulence regulation of Vibrio cholerae at the interface of host-gut microbiome interactions. Virulence. 2020;11(1):1582–1599.

38. Zhao W, Caro F, Robins W, Mekalanos JJ. Antagonism toward the intestinal microbiota and its effect on Vibrio cholerae virulence. Science. 2018;359(6372):210–213.

39. Schäfer J, Opgen-Rhein R, Strimmer K. Reverse engineering genetic networks using the “GeneNet” package. R News. 2006;6/5:50–53.

40. van Aert RCM. Meta-analyzing partial correlation coefficients using Fisher’s z transformation. Res Synth Methods. 2023;14(5):768–773.

41. Gene Ontology Consortium. Gene Ontology Consortium: going forward. Nucleic Acids Res. 2015;43(Database issue):D1049–D1056.

42. Kanehisa M, Goto S. KEGG: kyoto encyclopedia of genes and genomes. Nucleic Acids Res. 2000;28(1):27–30.

43. Schwechheimer C, Hebert K, Tripathi S, et al. A tyrosine phosphoregulatory system controls exopolysaccharide biosynthesis and biofilm formation in Vibrio cholerae. PLoS Pathog. 2020;16(8):e1008745.

44. Fong JNC, Yildiz FH. Biofilm Matrix Proteins. Microbiol Spectr. 2015;3(2):10.1128/microbiolspec.MB-0004-2014.

45. Dong TG, Mekalanos JJ. Characterization of the RpoN regulon reveals differential regulation of T6SS and new flagellar operons in Vibrio cholerae O37 strain V52. Nucleic Acids Res. 2012;40(16):7766–7775.

46. Götz F. *Staphylococcus* and biofilms. Mol Microbiol. 2002 Mar;43(6):1367–78.

47. Bi H, Deng R, Liu Y. Linezolid decreases *Staphylococcus aureus* biofilm formation by affecting the IcaA and IcaB proteins. Acta Microbiol Immunol Hung. 2022.

48. Zhou YH, Xu CG, Yang YB, et al. Histidine Metabolism and IGPD Play a Key Role in Cefquinome Inhibiting Biofilm Formation of *Staphylococcus xylosus*. Front Microbiol. 2018;9:665.

49. Pozzi C, Waters EM, Rudkin JK, et al. Methicillin resistance alters the biofilm phenotype and attenuates virulence in *Staphylococcus aureus* device-associated infections. PLoS Pathog. 2012;8(4):e1002626.

50. Zhang Z, Pan Y, Hussain W, Chen G, Li E. BBSdb, an open resource for bacterial biofilm-associated proteins. Front Cell Infect Microbiol. 2024;14:1428784.

51. Ishii N. GroEL and the GroEL-GroES Complex. Subcell Biochem. 2017;83:483–504.

52. Lin JS, Lai EM. Protein-Protein Interactions: Co-Immunoprecipitation. Methods Mol Biol. 2017;1615:211–219.

53. Rual JF, Venkatesan K, Hao T, et al. Towards a proteome-scale map of the human protein-protein interaction network. Nature. 2005;437(7062):1173-1178.

54. Button KS, Ioannidis JP, Mokrysz C, et al. Power failure: why small sample size undermines the reliability of neuroscience. Nat Rev Neurosci. 2013;14(5):365–376.

55. Tanay A, Regev A. Scaling single-cell genomics from phenomenology to mechanism. Nature. 2017;541(7637):331-338.

56. Saliba AE, Westermann AJ, Gorski SA, Vogel J. Single-cell RNA-seq: advances and future challenges. Nucleic Acids Res. 2014;42(14):8845–8860.

57. SoRelle ED, Dai J, Bonglack EN, et al. Single-cell RNA-seq reveals transcriptomic heterogeneity mediated by host-pathogen dynamics in lymphoblastoid cell lines. Elife. 2021;10:e62586.

58. Stegle O, Teichmann SA, Marioni JC. Computational and analytical challenges in single-cell transcriptomics. Nat Rev Genet. 2015;16(3):133–145.

59. Buettner F, Natarajan KN, Casale FP, et al. Computational analysis of cell-to-cell heterogeneity in single-cell RNA-sequencing data reveals hidden subpopulations of cells. Nat Biotechnol. 2015;33(2):155–160.

60. Mortazavi A, Williams BA, McCue K, Schaeffer L, Wold B. Mapping and quantifying mammalian transcriptomes by RNA-Seq. Nat Methods. 2008;5(7):621–628.

61. Shannon P, Markiel A, Ozier O, et al. Cytoscape: a software environment for integrated models of biomolecular interaction networks. Genome Res. 2003;13(11):2498–2504.

